# Estimating drivers of breast tissue transitions from normal to tumor state

**DOI:** 10.1101/2025.04.14.648655

**Authors:** Swapnil Kumar, Vaibhav Vindal

**Author notes:** Corresponding author: Prof. Vaibhav Vindal, Department of Biotechnology & Bioinformatics, School of Life Sciences, University of Hyderabad, Hyderabad, 500046, India., Email ID.

## Abstract

Tumor tissues are characterized by dysregulated gene expression patterns leading to altered cellular pathways and molecular functions as a result of their transition from normal to tumor state. Further, tumor-adjacent normal tissues (TANTs), utilized as a control in cancer research, are not molecularly normal and differ from healthy normal tissues. These TANTs represent a distinctive transitional state betwixt normal and tumor states. However, the mechanism underlying this state transition, expression dysregulation, and perturbed regulation remain largely unexplored and hence elusive. Herein, the transitions of breast tissues from normal and TANT to tumor states were modeled using gene expression and regulation data to estimate key drivers underlying these transitions. As a result, we identified 645 shared driver genes underlying the transitions of breast tissues from the healthy normal state to the adjacent normal and tumor states. Besides, we identified 635 shared driver genes underlying the transitions of TANTs to different subtypes. When we intersected both lists of shared driver genes, a total of 615 commonly shared driver genes across the state transitions were observed. Subsequently, functional annotations of these driver genes revealed their involvement in the growth and maintenance-related activity of cells. Additionally, key pathways associated with cancer pathogenesis, such as Wnt signaling, Notch signaling, NF-kappa B signaling, and PD-L1 expression and PD-1 checkpoint pathway in cancer, were found significantly enriched with these shared driver genes. Thus, the shared driver genes identified across tissue transitions provide ways forward to devise more efficient diagnostic and therapeutic strategies for early and effective disease management.

## Introduction

Breast cancer is a major women’s health concern due to high morbidity and mortality across the globe (Bray et al., 2024). It arises from abnormal growth in breast tissues, mainly lobules and ducts. It is complex and heterogeneous in nature and has multiple subtypes. These molecular subtypes are classified by the presence and absence of hormone receptors, viz., “Estrogen receptor” (ER), “Progesterone receptor” (PR), and “Human epidermal growth factor receptor 2” (HER2). Further, these subtypes of breast cancer include “Basal-Like” (Basal) (ER-/PR- and HER2-), “Luminal A” (LumA) (ER+/PR+, HER2-, and Ki67 low), “Luminal B” (LumB) (ER+/PR+, HER2+ or HER2-, and Ki67 high), “HER2-enriched” (Her2) (ER+/PR+ and HER2+), and “Normal-Like” (NormL) (ER+/PR+, HER2-, and Ki67 low) (Dai et al., 2015). Each of these subtypes exhibits different gene expression profiles and, hence, distinct gene regulatory network (GRN) systems. Besides, the gene expression profiles and GRNs of these subtypes also differ from their adjacent normal and even completely normal healthy tissues (Kumar & Vindal, 2023). Moreover, every cancer study employing patient samples include the tumor-adjacent normal tissue (TANT) samples as control for studying the tumor tissues. However, these morphologically and histologically looking normal TANTs are not completely normal and are different from healthy normal tissues (HNTs) derived from non-tumor-bearing healthy individuals. For example, the TANTs have various tumorigenic features, including altered pathways (Aran et al., 2017) and gene expression profiles (Kumar & Vindal, 2023). Despite this, there is a substantial lack of related knowledge as well as thorough experimental studies on the driver genes underlying the transitions of normal breast tissues into different subtypes of the same breast cancer.

It is experimentally well-documented that alterations in gene expressions lead normal tissues to proceed toward diseased ones. When normal tissues transform into tumors, the exact attribute of the alteration of cellular pathways and loss of functions specifically depends on differentially expressed genes and pathways varying between different cell or tissue types. Therefore, specific states of any tissues, e.g., normal and tumor tissues, harbor distinct gene expression profiles. Further, these normal and tumor states might be representative of changes that occurred during the transition or transformation from one to another condition, e.g., healthy to tumor condition. Different tissue types can be classified by their gene expression profiles and, hence, GRNs; consequently, the transitions between different states can be modeled using any changes occurred in their underlying GRNs.

In this study, we have applied modeling of network state transition using gene expression and regulation data, a regression-based method implemented in MONSTER (Schlauch et al., 2017), for inferring transcription factors (TFs) as drivers of tissue state transformation at the GRN level. We analyzed TANTs along with all five subtypes of breast cancer to identify TFs that lead to the alteration of the network structure as the healthy tissues progress toward the diseased state. For this, first, TF-target gene regulations were predicted using gene expression profiles of TANTs and subtypes; then, driver genes and TFs present in TANTs and subtypes were identified using the corresponding expression profiles coupled with the predicted TF-target gene regulations. Thus, our results demonstrate the key candidate genes and TFs associated with TANTs and subtypes, offering a promising avenue toward designing and developing more efficient, early-stage-specific diagnostic and therapeutic interventions.

## Methods

### Acquisition of tumor and normal tissue data

Gene expression datasets of tumor tissues and TANTs were retrieved using the recount2 project (Collado-Torres et al., 2017) from the portal of genomic data commons of “The Cancer Genome Atlas” (TCGA) (Tomczak et al. 2015; Weinstein et al. 2013). However, the gene expression data of healthy normal tissues (HNTs) were retrieved using the recount2 project from “The Genotype-Tissue Expression” (GTEx) portal (Lonsdale et al., 2013). These data were downloaded with the help of the “TCGAbiolinks” R package (Colaprico et al., 2016). Subsequently, all three types of gene expression data were pre-processed, filtered, and normalized using the “TCGAbiolinks” R package for further analysis.

### Motif mapping data for TF-target gene regulations

Motif mapping data contain information on which genes have likely binding sites for a particular TF in their promoter and vicinity. This serves as the prior on TF-gene regulations and informs the network inferential method with a prior network of TFs targeting genes. Each row of this prior data defines one regulatory edge with TF, the corresponding targeted gene, and the edge’s strength. For the current study, we have considered networks as unweighted and, hence, have populated the entire data with one as the strength of the edges. The TF-gene motif mapping data were collected from the motif mappings provided in online documentation and resources of PANDA (Glass et al., 2013).

### Prediction of transitioning drivers

Driver genes underlying the transition of breast tissues from the normal state to the tumor state were predicted using the MONSTER package (Schlauch et al., 2017) as implemented in netZooR v1.4 (Ben Guebila et al, 2023) and R v4.2. It first predicts the regulatory relations of TFs and their target genes using the PANDA implementation (Glass et al., 2013) by utilizing gene expression data of each tissue type along with prior information on the TF-gene motif mapping. Herein, the gene expression data of genes with counts per million (CPM) values of one or more across 50% or more samples were only considered in the case of each tissue type. However, the predicted gene regulations for initial and terminal conditions, i.e., normal and tumor tissues using the PANDA package, can also be used directly as input in the MONSTER algorithm for the transition driver predictions. Subsequently, this calculates a transition matrix by mapping the GRN of the initial state to the GRN of the final state and follows the identification of a set of significant and critical driver genes potentially responsible for the transition of normal tissues and TANTs to tumor tissues. To determine the statistical significance of the results produced by the MONSTER, the samples were permuted 100 times, and the analysis was re-run for each permutation. Further, the state transition matrices, along with the rewired regulations of differential TFs involved in the state transition process, were visualized.

### Functional and pathway annotation of key drivers

Key driver genes for the modeled tissue transitions were checked for their involvement in molecular functions (MFs) and biological processes (BPs) utilizing the Gene Ontology (GO) and pathways by the KEGG pathway-based over-representation analysis method implemented in the clusterProfiler R package (Xu et al., 2024). To identify significantly associated or enriched MF, BP, and pathway terms with key driver genes, the significance level was set to an adjusted P-value of 0.05. Herein, the P-value was adjusted by the method of “Benjamini-Hochberg” (BH) to control the false positive rate.

### Disease association of key drivers

Driver genes underlying the transition of tissues from a normal to tumor state were subjected to Disease Ontology (DO)-based enrichment analysis using the DOSE R package (Yu et al., 2015) with the “BH” method of P-value adjustment. All the enriched terms with adjusted P-values ≤ 0.05 were regarded as significant for the result inference purpose. Further, these driver genes were also searched for their previously known association with any diseases, including cancers, using disease-gene association information from the database of DisGeNET (Piñero et al., 2020) and the catalog of known and candidate cancer genes from the database of Network of Cancer Genes (NCG) (Repana et al., 2019).

## Results and discussion

### Expression data of tumor tissues, TANTs, and HNTs

Gene expression profiles of all six tissue types were processed together for further analysis. Herein, only expressions of protein-coding genes were considered as per the list of genes and their details retrieved from the Human Gene Nomenclature Committee. The detailed distribution of samples and genes across all tissue data are provided in Table 1. The LumA subtype had the highest number of samples (570), while the NormL subtype had the lowest (40). The gene expression data of only female samples, from both TCGA and GTEx, were used for the current study.

**Table 1.** Samples and genes across all six tissue types.

| Tissue | Samples | Genes | PC Genes |
| --- | --- | --- | --- |
| Basal | 200 | 58037 | 19079 |
| Her2 | 82 | 58037 | 19079 |
| LumA | 570 | 58037 | 19079 |
| LumB | 207 | 58037 | 19079 |
| NormL | 40 | 58037 | 19079 |
| TANT | 111 | 58037 | 19079 |
| HNT | 92 | 58037 | 19079 |

PC Genes, Protein-coding genes.

### Driver genes underlying transitions of tumors and TANTs from HNTs

The modeling of state transition from normal state, i.e., HNTs, to tumor state, i.e., different subtypes and TANTs, resulted in the identification of differential TFs as potential drivers involved in the state transitions. Herein, we have considered the TANTs similar to tumors because it is well-proven that the TANTs are not entirely normal and have tumorigenic features at the molecular level. When state transitions were modeled for HNTs with tumor tissues and TANTs, a maximum of 830 driver genes were identified for Basal, Her2, and TANT, whereas a minimum of 812 for LumA. The detailed distributions of identified driver genes across all modeled transitions are available in Table 2 and Fig. 1. There were a total of 645 commonly shared driver genes underlying the modeled transitions of tumor tissues and TANTs from the HNTs (Table S1). These shared common driver genes included some well-known cancer-associated genes such as *ATM, BCL11A, BRCA2, E2F1, ESR1, FOXA1, JUN, MYC, RUNX1, TP53*, and *TP63*.

**Table 2.** Distribution of genes with a CPM value of more than one and number of drivers identified across all possible transitions.

| Transition | | PC Genes (CPM ≥ 1) | Drivers |
| --- | --- | --- | --- |
| From | To |  |  |
| HNT | Basal | 14233 | 830 |
|  | Her2 | 14362 | 830 |
|  | LumA | 14127 | 812 |
|  | LumB | 14174 | 815 |
|  | NormL | 14398 | 821 |
|  | TANT | 14640 | 830 |
| TANT | Basal | 14504 | 850 |
|  | Her2 | 14534 | 828 |
|  | LumA | 14196 | 820 |
|  | LumB | 14463 | 839 |
|  | NormL | 14270 | 808 |

PC Genes, Protein-coding genes; CPM, Counts per million.

**Fig. 1.**
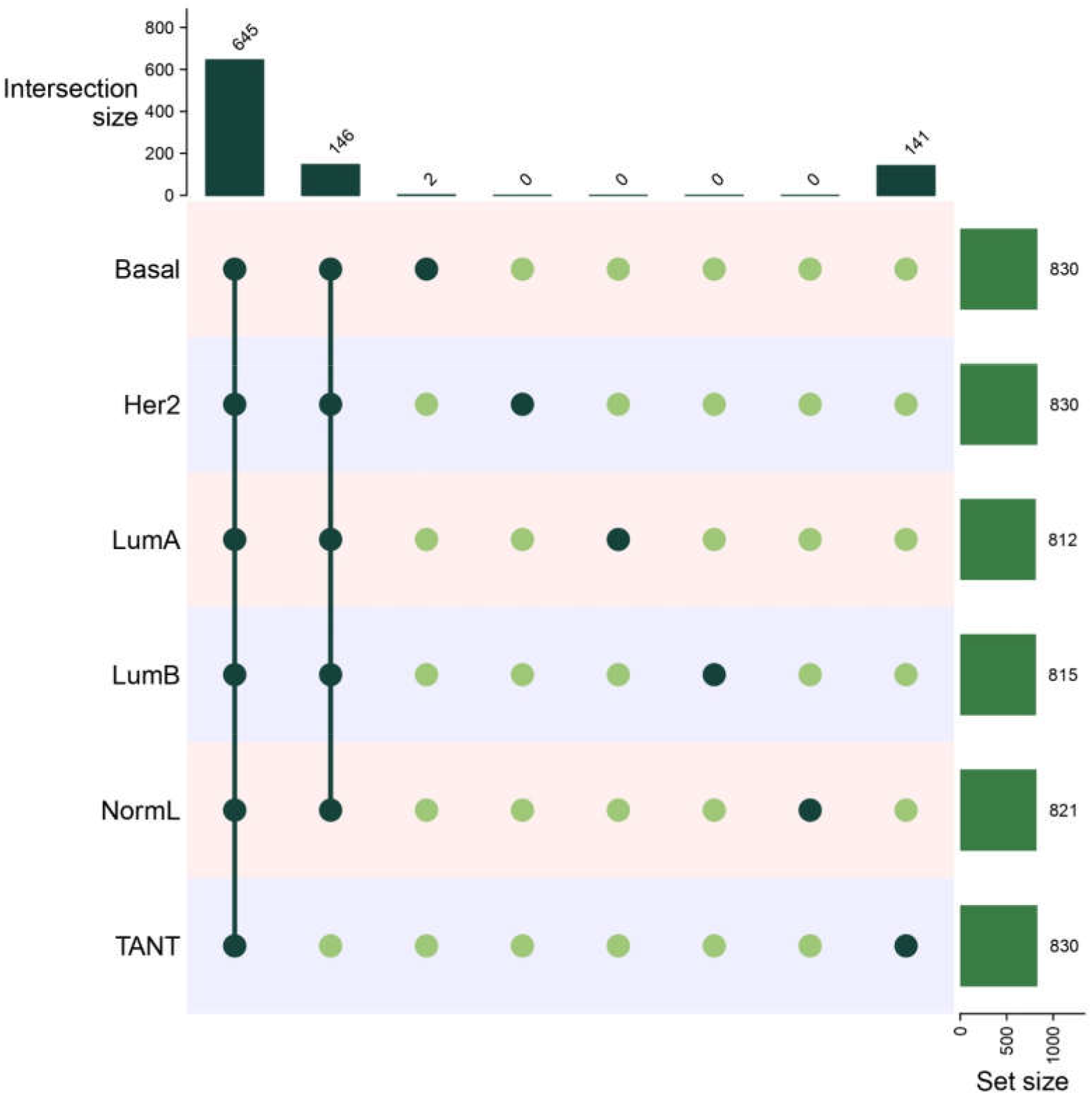
Distribution of driver genes underlying transitions of tumor tissues and TANTs from HNTs.

### Driver genes underlying transitions of subtypes from TANTs

We also modeled the state transitions of tumor tissues (five subtypes, viz., Basal, LumA, LumB, Her2, and NormL) from the TANTs, which resulted in the identification of differential TFs as potential drivers involved in the respective state transitions. Consequently, a maximum of 850 driver genes were identified for the Basal subtype, whereas a minimum of 808 for the NormL. The detailed distributions of identified driver genes across all modeled transitions are available in Table 2 and Fig. 2. There were a total of 635 commonly shared driver genes across all the five modeled state transitions of tumor tissues (breast cancer subtypes) from the TANTs (Table S2). These shared common driver genes across transitions also included some well-known cancer-associated genes, such as *ATM, BCL11A, BRCA2, E2F1, ESR1, FOXA1, JUN, MYC, RUNX1, TP53*, and *TP63*, which were even present in the transitions of HNTs to tumor tissues.

**Fig. 2.**
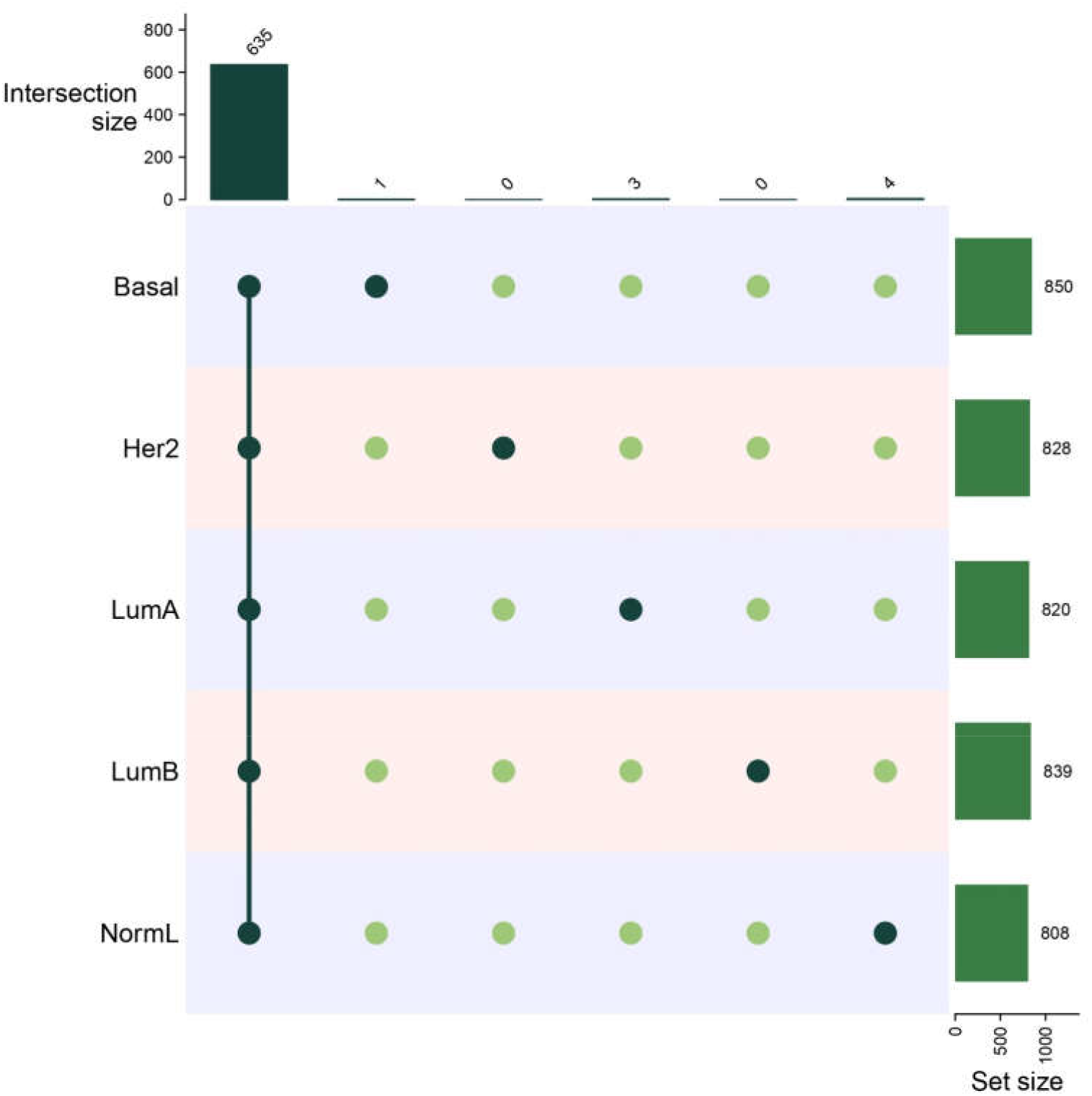
Distribution of driver genes underlying transitions of tumor tissues from TANTs.

When we looked for the common driver genes in both transitions, i.e., from HNTs to TANTs and tumors and from TANTs to tumors, most of the driver genes from both transitions were common, and hence, there were 615 shared driver genes. These commonly shared driver genes across all the transitions can serve as key candidates for the design and development of potential diagnostics and therapeutics specifically for the effective management and treatment of the disease at an early-stage. It has been reported that the mutated *ATM* gene is associated with a higher breast cancer risk and contributes to its development (Dork et al., 2001; Soukupova et al., 2008; Bogdanova et al., 2009). Next, the gene *BCL11A*, a B-cell proto-oncogene, has previously been found to be associated with transcriptional regulation and as a mutated gene in breast cancer (Sjoblom et al. 2006). Similarly, the DNA repair-associated gene *BRCA2* has been identified as a mutated gene in the case of breast cancer (Valarmathi et al., 2004; Hall et al., 2009; Vaidyanathan et al., 2009).

### Functions and processes of key drivers

GO-based enrichment analysis revealed a total of 66 MF and 369 BP terms significantly enriched or over-represented with commonly shared driver genes. Among all enriched MF terms, terms related to transcription regulations, including “DNA-binding transcription activator and repressor activity,” “transcription coactivator and corepressor activity,” “histone deacetylase activity,” and “promoter-specific chromatin binding” were most significantly enriched with the maximum number of the shared driver genes (Fig. 3). Moreover, among all enriched BP terms, terms related to developmental and transcriptional processes, such as “gland development,” “mononuclear cell differentiation,” “intracellular receptor signaling pathway,” “regulation of miRNA transcription,” “miRNA metabolic process,” etc., were the most enriched terms with the majority of the shared driver genes (Fig. 4). These MF and BP terms associated with the shared driver genes indicate that these driver genes play vital roles in the growth and maintenance-related activity of cells and, hence, the constituting tissues.

**Fig. 3.**
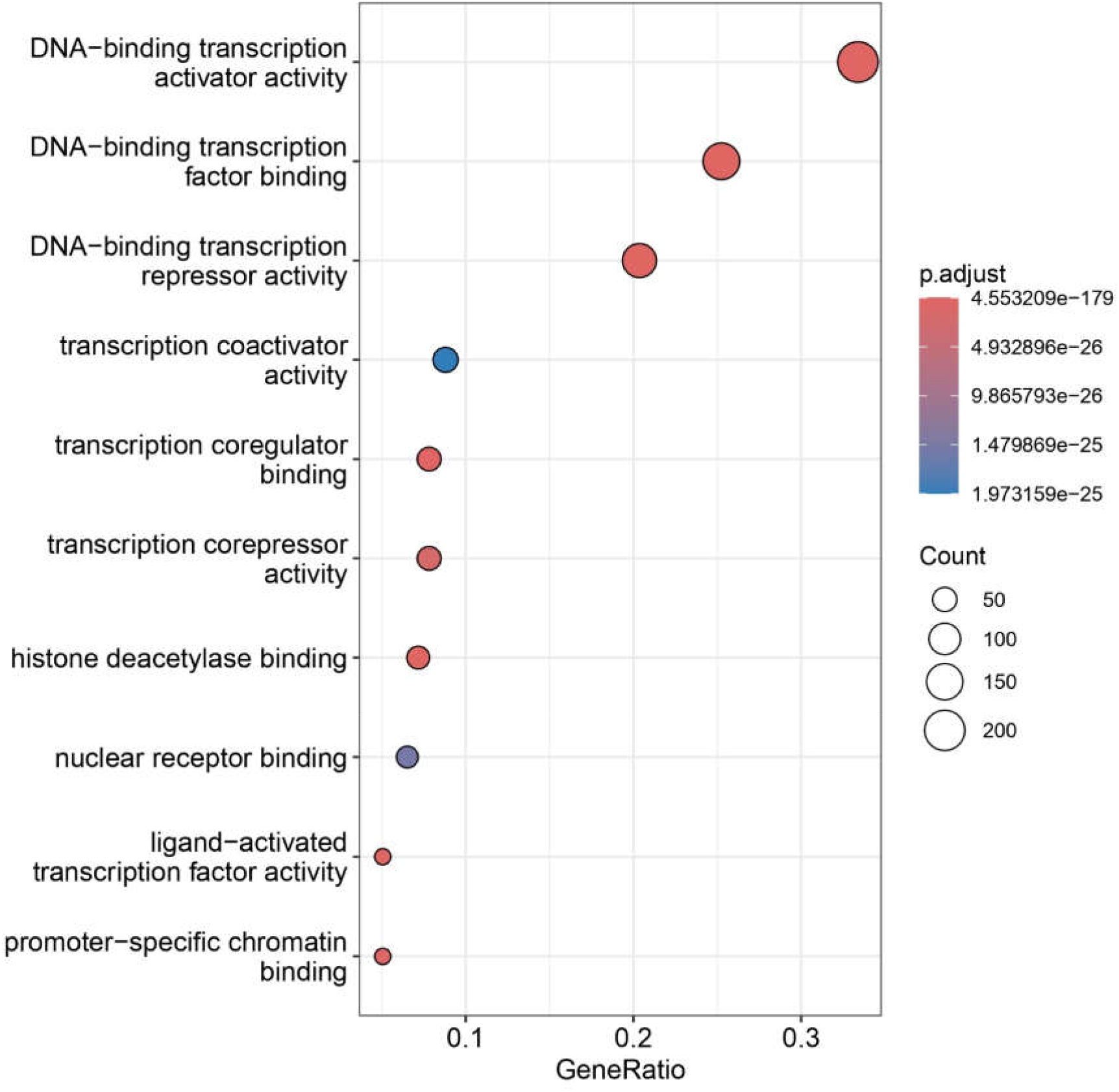
Top MF terms significantly associated with shared driver genes.

**Fig. 4.**
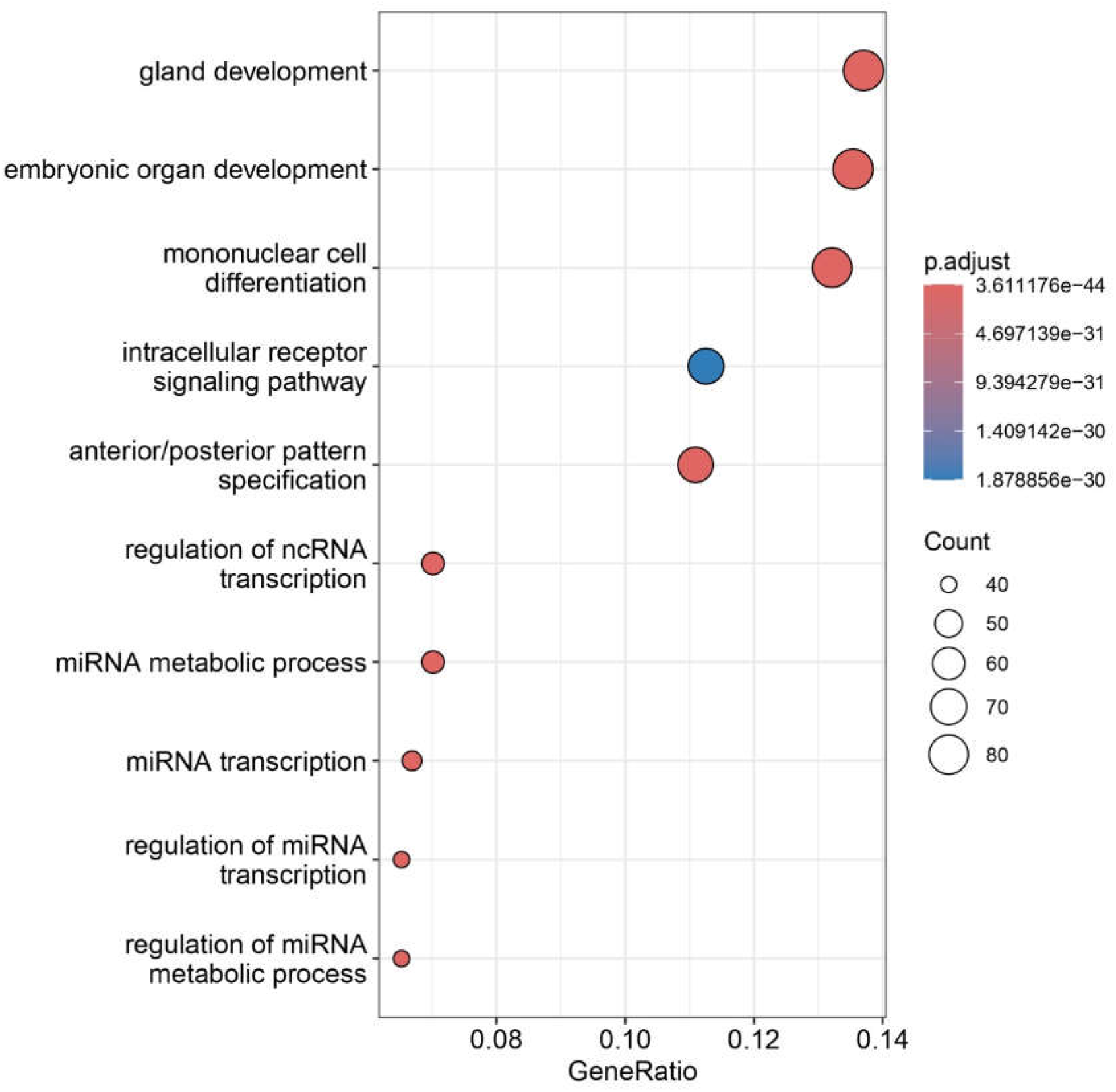
Top BP ontology terms significantly associated with shared driver genes.

### Enriched pathways of key drivers

When the commonly shared driver genes across all types of modeled transitions were subjected to pathway analysis, 83 pathways were identified as significantly associated with these driver genes. Among all these pathways, the pathway of “Transcriptional misregulation in cancer” (hsa05202) was found to be significantly enriched or over-represented with the maximum number of shared driver genes. Transcriptional misregulation is a hallmark of cancer that is driven by genetic alterations that disrupt the transcriptional control of genes. Further, genetic alterations can affect transcriptional controls at many levels, including Cis-elements and Trans-factors, which can lead to a variety of alterations in cellular properties that contribute to tumor development. These alterations in cellular properties can include dysregulated transcriptional programs (Koschmieder et al., 2009) and fusion oncoproteins (Prensner & Chinnaiyan, 2009).

Other significantly enriched pathways included “Breast cancer pathway,” “TGF-beta signaling pathway,” “Notch signaling pathway,” “Wnt signaling pathway,” “Cell cycle,” “Estrogen signaling pathway,” “MAPK signaling pathway,” “NF-kappa B signaling pathway,” “Toll-like receptor signaling pathway,” and “PD-L1 expression and PD-1 checkpoint pathway in cancer” (Fig. 5). It is interesting to note that these pathways are well-known for their involvement in the pathogenesis of various types of cancers, including breast cancer. For example, the altered form of the MAPK signaling pathway is known for its association with the development of various diseases, including cancers (Qi & Elion, 2005; Kim & Choi, 2010). Thus, these significantly over-represented pathways with shared driver genes highlight their importance in the transitions of normal tissues to tumor tissues and, hence, the tumorigenesis of breast cancer.

**Fig. 5.**
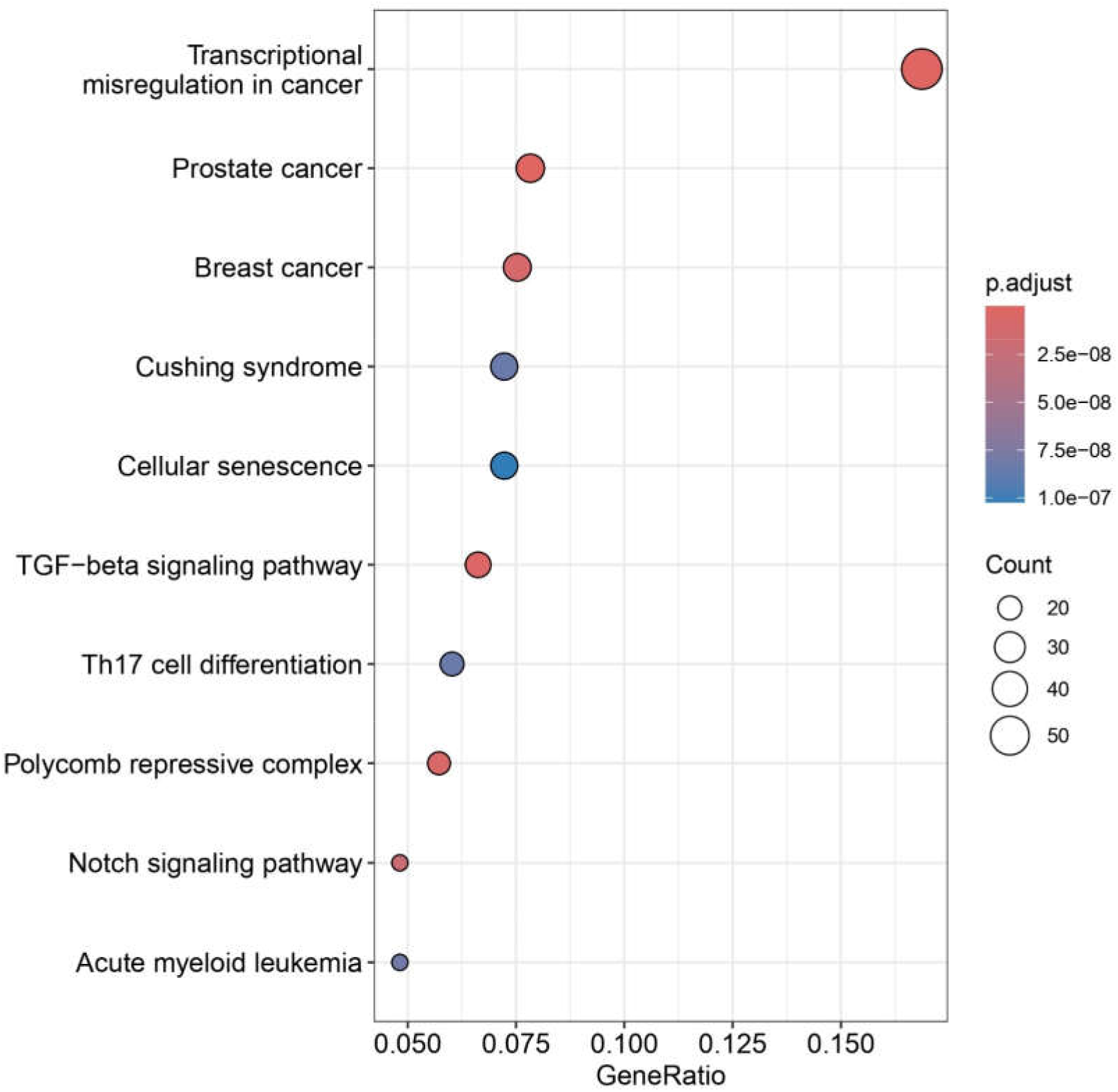
Top KEGG pathway terms significantly associated with shared driver genes.

### Enriched diseases of key drivers

The DO and NCG-based enrichment analysis of the shared driver genes revealed their significant associations with various cancer types and other diseases. It was noticed that 135 DO terms and 37 NCG terms were significantly enriched with these common driver genes. Among DO terms, cancer-related terms such as “breast cancer,” “lung cancer,” “liver cancer,” and “prostate cancer” were enriched with a majority of the shared driver genes (Fig. 6). Similarly, NCG terms, including “breast cancer,” “bladder cancer,” “liver cancer,” and “prostate cancer,” were found to be significantly enriched with the majority of the shared driver genes (Fig. 7). These enriched terms show that all the shared driver genes identified herein have potential implications in the transitions of normal breast tissues to tumor forms.

**Fig. 6.**
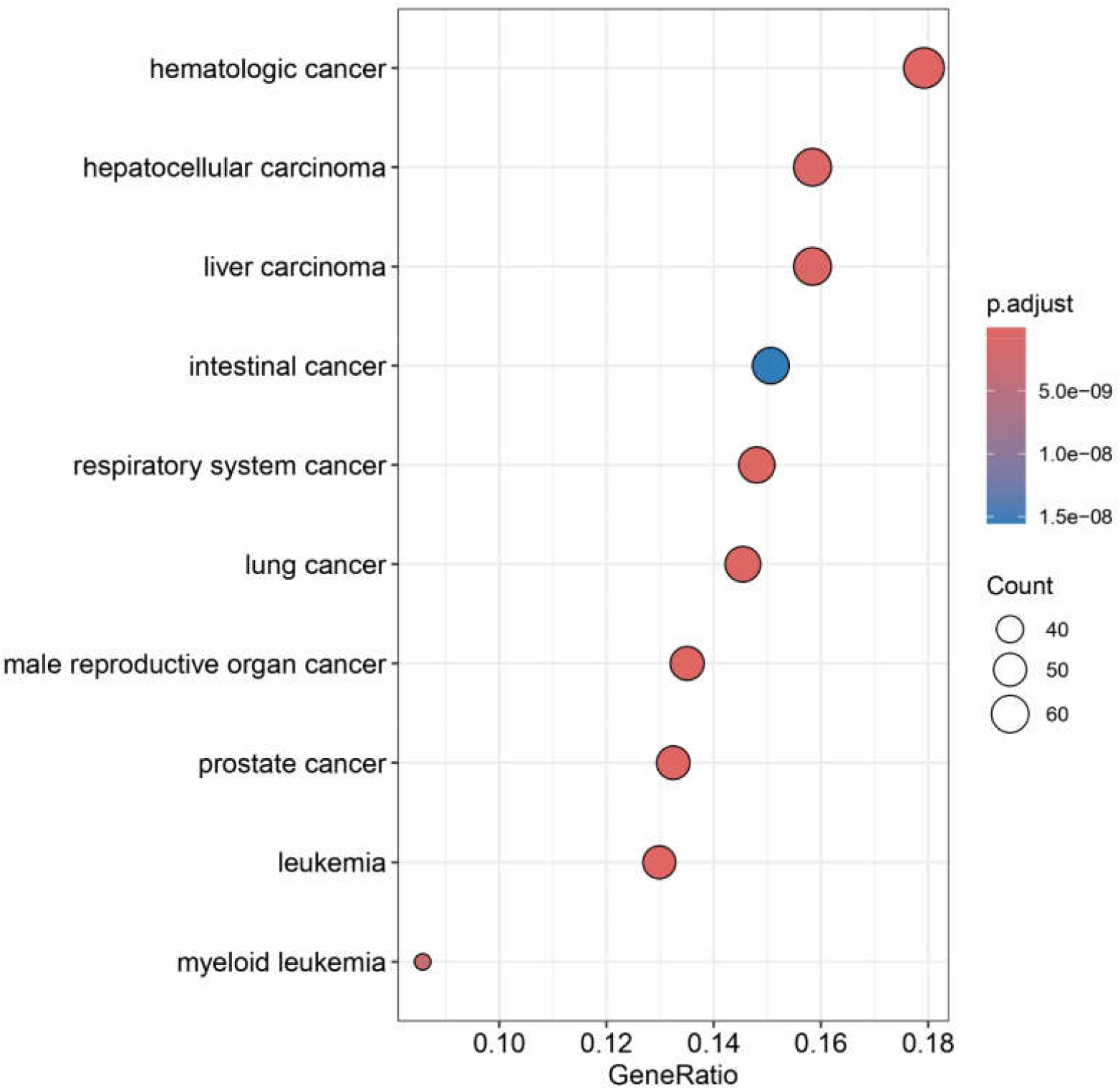
Disease ontology terms significantly associated with shared driver genes.

**Fig. 7.**
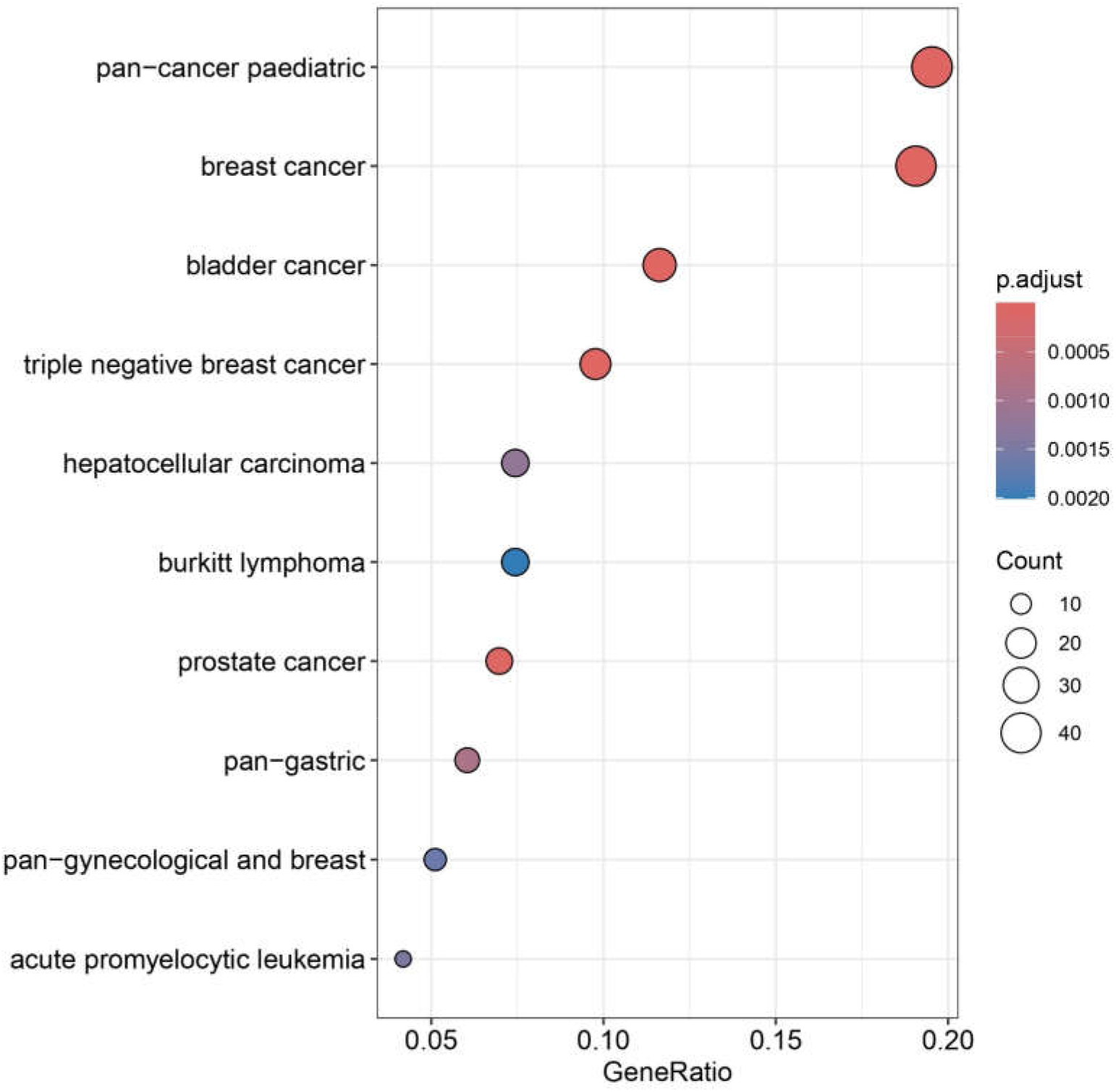
NCG terms significantly associated with shared driver genes.

### Limitations of the current study

It is the first study to date on modeling the transitions of breast tissues from normal to tumor state using gene expression and TF-gene regulatory data of tumor tissues, TANTs, and normal tissues to estimate the underlying driver genes. However, the present study has some limitations, too. First, we have considered the samples of TANTs as one group of tissues that represents the intermediate state between normal and tumor states, irrespective of which subtypes these TANTs belong to due to the low number of samples of TANTs to individual subtypes. Thus, with the future availability of more expression data on TANTs of different subtypes, transitions of each subtype can be modeled to estimate the underlying driver genes towards the inference of pathway mechanisms transforming breast tissues from normal to tumor forms more accurately and efficiently. Second, we computed only 10 null permutations to determine the statistical significance of our result for each modeled transition due to the requirement of a large amount of computational processing therein. However, computing high numbers of null permutations, such as 1000, would likely ensure more sufficient null observations for estimating null distributions and computing empirical P-values. In turn, it will produce comparatively more significant predictions of transitioning driver genes. Third, annotations retrieved from the databases of GO and pathways tend to be biased toward genes and pathways that are well-studied and properly characterized. Consequently, genes that lack functional and pathway-based annotations get ignored while performing GO and pathway-based annotation analyses; hence, those genes are required to be investigated separately.

## Conclusions

The transitions between two states of phenotypes, viz., normal and tumor, can be modeled in terms of gene expression space, where a change in phenotype correlates with the transition between gene expression profiles of the two states. Here, we implemented the framework of network modeling for inferring state transitions between HNT/TANT and five subtypes (Basal, LumA, LumB, Her2, and NormL) using gene expression and regulatory data by assuming that gene expression patterns were driven by perturbations in gene regulation patterns and therefore, subsequent changes in the underlying GRN. It helped us to explore and decode how phenotypic alterations and the associated regulatory changes were driven by the “rewiring” of TF-target regulations and to identify all those TFs that were involved in the alteration of their targets. As a result, it revealed critical driver genes underlying the transitions of all five subtypes from the HNTs and TANTs and also the TANTs from the HNTs. These key drivers included genes such as *ATM, BCL11A, BRCA2, E2F1, ESR1, FOXA1, JUN, MYC, RUNX1, TP53*, and *TP63*. Thus, the current study paved the way to identify and explore the TFs that drive the transition occurring from normal healthy to tumor states, and it can be applied to a wide range of systems, such as Type 2 diabetes, Cardiovascular diseases, Neurodegenerative diseases, and other types of cancers.

## Acknowledgements

The authors thank the Center for Modeling, Simulation & Design (CMSD), University of Hyderabad, for providing computational facilities. Further, VV would like to thank the Indian Council of Medical Research (ICMR), New Delhi (ISRM/12(72)/2020, ID: 2020-2951), Institution of Eminence – University of Hyderabad (No. UoH/IoE/RC3-21-052), and Department of Biotechnology (DBT), Government of India (No. BUILDER-DBT-BT/INF/22/SP41176/2020), for their financial support. SK also acknowledges the Institution of Eminence – UoH for the Performance-Based Publication Incentives, University of Hyderabad for the Non-NET Fellowship, and ICMR for the Senior Research Fellowship (Grant No.: 3/2/2/113/2019/NCD-III, ID: 2019-6723).

## Funding

This study was supported by the extramural research grant from the Indian Council of Medical Research (ICMR) New Delhi (ISRM/12(72)/2020, ID: 2020-2951).

### Contributions

SK and VV conceived and designed the study. SK performed the experiments and analyzed the data. SK drafted, wrote, and edited the manuscript. VV reviewed and edited the manuscript. VV supervised the study. All authors read and approved the final manuscript.

### Ethics Statement

This manuscript does not involve human subjects, animals, or any form of experimentation requiring ethical approval or consent.

## Data availability statement

The common driver genes for two transitions, viz., transition from HNTs to TANTs and tumor tissues and transition from TANTs to tumor tissues identified in the current study are available in the supplementary data files. Further, the gene expression datasets of tumors, TANTs, and HNTs used in this study are publicly available and can be downloaded from the portals of TCGA (tumors and TANTs) and GTEx (HNTs).

## Supplementary Tables

Table S1. Driver genes underlying transition from HNTs to TANTs and tumor tissues

Table S2. Driver genes underlying transition from TANTs to tumor tissues

